# Deep long-read metagenomic sequencing reveals niche differentiation in carbon cycling potential between benthic and planktonic microbial populations

**DOI:** 10.1101/2024.06.04.597336

**Authors:** Tomeu Viver, Katrin Knittel, Rudolf Amann, Luis H. Orellana

**Affiliations:** Department of Molecular Ecology, Max Planck Institute for Marine Microbiology, Bremen, Germany

**Keywords:** metagenomics, sediment microbial communities, Helgoland, polysaccharide utilization loci, chemolithoautotrophy, long-read sequencing

## Abstract

Coastal marine sediments function as large-scale natural biocatalytic filters, remineralizing and transforming organic matter. Benthic microbiomes exhibit remarkable temporal stability, contrasting with the dynamic, substrate-driven successions of bacterioplankton. Nonetheless, understanding their role in carbon cycling and interactions between these microbial groups is limited due to the complexity of benthic microbiomes. Here, we used a seasonally resolved, deep short- and long-read metagenomic approach to examine distinctive genomic features of microbiomes recovered from sediment, the overlaying water column, and particle-attached bacteria and archaea in the North Sea. We recovered 115 benthic metagenome-assembled genomes (MAGs) that belonged to *Woeseiales*, *Rhizobiales*, *Planctomycetia*, *Gemmatimonadota*, and *Desulfobacterota* species. While *Proteobacteria* and *Actinobacteriota* were characteristic phyla of sediments, *Acidimicrobiia* and *Desulfocapsaceae* species were shared between sediments and particle-attached fractions of the water column, indicative of significant bentho-pelagic coupling. Predominant members of the family *Woeseiaceae* carried polysaccharide utilization loci (PULs) predicted to target laminarin, alginate, and α-glucan in sediments. In contrast, water column species lacked PULs and encoded a significantly higher fraction of sulfatases and peptidases, indicating a role in the degradation of protein-rich and sulfated organic matter. Our findings disentangle family-level adaptations and niche differentiation between globally significant benthic and water column populations involved in marine organic matter degradation and carbon storage.

## Introduction

Gaining insight into the ecological niches that allow the colonization, proliferation, and establishment of microbial communities in a given environment remains a fundamental challenge in microbial ecology^1^. A comprehensive understanding of the environmental conditions (abiotic factors), the biotic interactions among organisms, and the diverse microbial metabolic pathways are imperative to delineate microbial ecological niches. The niche theory has mainly gained significance in elucidating the adaptability of microorganisms within marine environments, specifically concerning their responses to seasonal variations^2^, that for the most part are driven by environmental factors, such as temperature, salinity^3,4^ and substrate availability^5,6,7^. Nonetheless, the metabolic differentiation of taxonomically related microorganisms inhabiting marine sediments compared to those in pelagic environments, characterized by differences in substrate availability, has remained relatively understudied.

While bacterioplankton exhibited considerable variability and dynamics in microbial populations, sediment-dwelling microbial communities demonstrate remarkable temporal stability^8,9,10^. For instance, in the North Sea, phytoplankton spring blooms have been shown to cause substrate-based successions of pelagic taxa such as *Polaribacter, Roseobacter*, *Flavobacteria*, *Verrucomicrobia*, SAR92 and archaeal populations^2,6,11,12,13,14^. In contrast, sediment communities exhibit a much higher stability even in the presence of phytodetrital inputs, featuring dominant taxa like *Woeseiaceae*, *Desulfobacterota*, and *Planctomycetota*^9,10^.

Members of the *Woeseiaceae* family have been recognized as major parts of the benthic microbial communities dominating in various settings, ranging from shallow-sea coastal sediments^15,16,17,18,19^ to deep ocean seafloor^20,21,22^, hydrothermal vent chimneys^23^, as well as in both, oxic and anoxic environments^24^. Ecological and phylogenetic studies have described the *Woeseiaceae* family as a widely distributed group within the class *Gammaproteobacteria*, exhibiting significant prevalence in marine sediments worldwide^21^. Moreover, species within the *Woeseiaceae* family have been characterized by their versatility in metabolic strategies, encompassing both heterotrophic and chemolithotrophic metabolisms^15,21,25^. The potential for chemolithotrophy, involving sulfide and hydrogen oxidation, has been well-documented. Their heterotrophic metabolism has been linked to the utilization of mono- and polysaccharides and, proteolytic activity, indicating a significant role in mediating the turnover of protein-rich and sulfated organic matter, including remnants of cell membranes or walls^15,21,25^. Additionally, *Woeseiaceae* species have been implicated in nitrogen cycling and the release of N_2_O from sediments^21,26^. These diverse metabolic pathways suggest a high ecological diversification among species of the *Woeseiaceae*. Yet, due to a lack of cultivability and in the absence of a high quality genome information, the genetic bases of the niche differentiation and their implications for organic matter recycling remain limited.

Here, we applied a deep long-read (LR) metagenomic approach to explore the diversity and metabolic capabilities of the sediment microbial communities in the upper layers of aerobic coastal sandy sediment samples from Helgoland, over a year-long temporal series in 2018. LR metagenomics offers the advantage of recovering full-length 16S rRNA gene sequences, allowing accurate taxonomic analyses of the complete sediment microbial community^27^. Moreover, we compared sediment microbial diversity with those from the overlaying water column during the phytoplankton spring bloom, including free-living and particle-attached bacteria. The recovery of high-quality *Woeseiaceae* MAGs from sediments and overlaying water column samples allowed us to detect niche-specific and metabolic potential capabilities at the species level.

## Results

### Metagenome description and microbial diversity

With five metagenomes we covered the sampling period from March 7^th^, 2018 to January 18^th^, 2019. For evaluating of the effect of sequencing depth, five replicates were sequence from the initial sample (Mar18_A1 to A5). The average output for each metagenomic sample was 30.4 Gbp [interquartile range (IQR): 27-31 Gbp] and average read length of 5.1 kbp (IQR: 3.2-6.5 kbp) (Table 1 and S1). Thus, a total of 274 Gb of long-read (LR) PacBio HiFi sequencing data were generated. We observed temporal stability of the sediment microbial populations when using a beta diversity index based on k-mer distances, with values ranging from 0.063 to 0.085 (on a scale ranging from 0 (no differences) to 1 (completely different); (Figure S1 and Table S2). The distance values among the Mar18 replicates were lower (average 0.067 ± 0.009) when compared with the other sampling points (average 0.08 ± 0.003). Based on the taxonomic affiliation of unassembled LRs, most of the sequenced metagenomic samples corresponded to bacterial sequences, and the proportion assigned to eukaryotes ranged between 6.3% and 6.9% (Table S1).

**Table 1:**
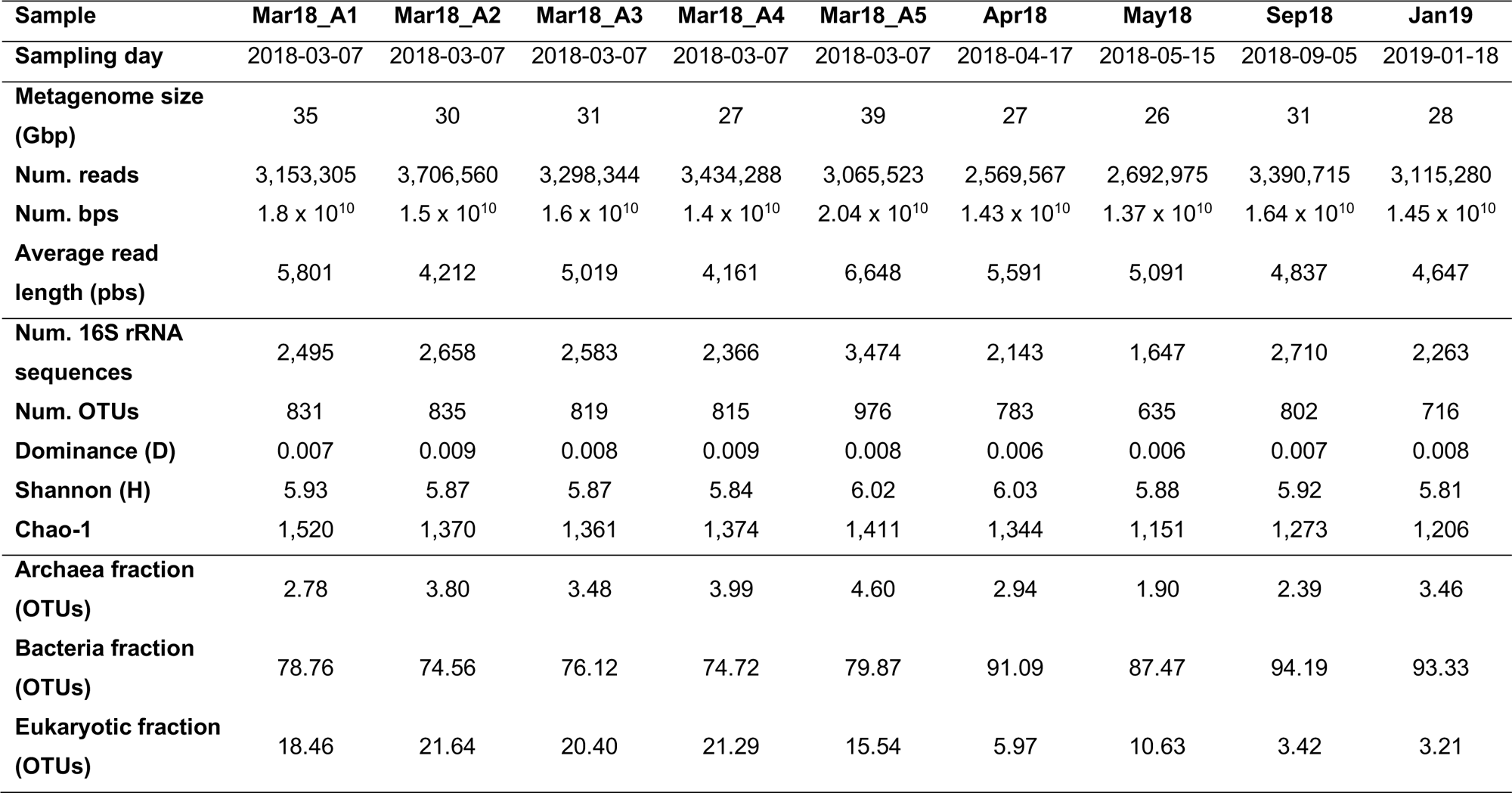
Long-Read metagenomic statistics of Helgoland sediment samples.

To obtain a broad overview of the community composition, we first exploited the recovery of full-length SSU rRNA gene sequences from all unassembled LRs. A total of 26,082 sequences (average 1,457 ± 143 bp length) were extracted from unassembled LRs, and after clustering the sequences at 98.7% identity, 1,732 OTUs were identified (Table 1). Of these, 1,618 were affiliated to Bacteria, 8 to Archaea, and 106 to Eukaryota (Table S3). Despite the high sequencing effort, the non-saturated rarefaction curves of each metagenome individually indicated a low metagenomic community coverage of the bacterial populations (Figure S2A). Nevertheless, rarefaction reached saturation by combining all metagenomes (Figure S2B). The taxonomic classification of OTUs indicated that the most abundant classes were *Gammaproteobacteria* and *Alphaproteobacteria* (average abundance of 32 ± 2.6% and 18 ± 4.7%, respectively), followed by *Bacteroidia* (5.1 ± 1.3%), *Planctomycetes* (4.5 ± 0.5%), and *Acidimicrobiia* (4.3 ± 1.0%) (Figure S3 and Table S5). At the OTU level, the Archaea *Candidatus* Nitrosopumilus was the most abundant species, comprising 3.8 ± 1.2% of the microbial community. Interestingly, within the class *Gammaproteobacteria,* the family *Woeseiaceae* comprised the most diverse family (represented by 50 OTUs) and reached an abundance of 7.6 ± 0.9%, similar to what was previously observed using 16S rRNA sequence amplicons^9^ (Table S6). In the replicated Mar18 samples, eukaryote OTUs accounted for 19.5 ± 2.5% of the total community, while in all other samples, less than 10.6% (Table S1). The most abundant Eukaryotes were assigned to copepods, the worm *Monostilifera,* and the animal parasite *Acetosporea*. At the class level, similar relative abundances were determined based on 16S gene amplicons, full-length 16S extracted from long-reads, and unassembled total metagenomic reads methodologies (Figure S4). At the family and genera levels, the taxonomic distribution and relative abundance of 16S rRNA sequences from metagenomes were statistically similar to those previously observed using amplicon 16S rRNA amplicon sequencing^9^ (Figure S5).

### Diversity and novelty in long-read derived metagenome-assembled genomes (MAGs)

Due to the observed stability in the microbial composition of the sediment samples and to assess the effect of sequencing effort in the representation of species in MAGs, we employed two co-assembly strategies. When we concatenated the metagenomic files chronologically, we observed a gradual increment in the number of recovered MAGs (Figure 1). Starting with 8 MAGs from the initial single metagenome, we reached 75 MAGs after concatenating the five replicates from sample Mar18 (A1 to A5). We increased the number of recovered MAGs to 92 when the remaining four metagenomes (from Apr18 to Jan19) were included. Similarly, we observed a linear increase in MAGs when concatenating all samples and subsampling to match the Gbps size of the previous strategy. These results highlight the benefits of increasing the sequencing effort for the recovery of MAGs in complex samples when using LR metagenomic approaches (Figure 1).

**Figure 1:**
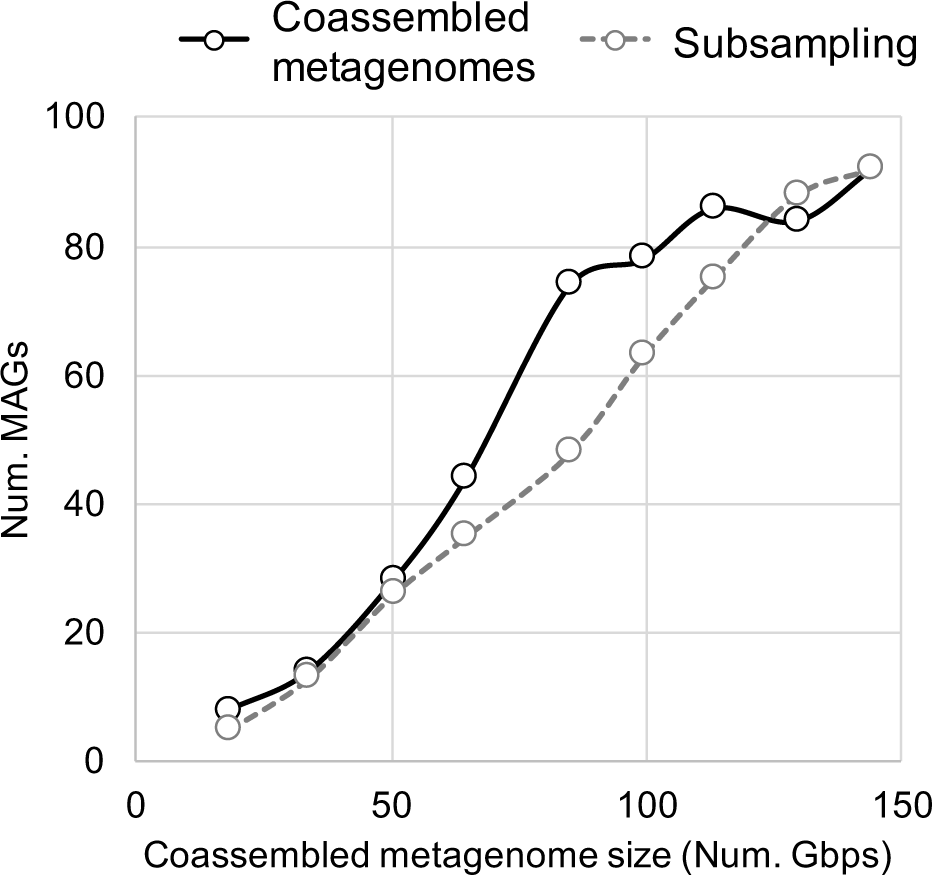
Co-assembled long-read metagenomes increase the recovery of medium and high-quality MAGs from sediment metagenomic samples. MAGs recovered after consecutive co-assembly of metagenomic samples, following the temporal scale order (represented in black line), and MAGs recovered through subsampling of all concatenated metagenomes (discontinuous grey line).

All MAGs recovered using concatenated metagenomic samples were de-replicated, resulting in 115 genomospecies (gspp) at 96% average nucleotide identity (ANI) value. Their genomic completeness ranged between 50% and 98.3% (average 79.2 ± 12%), and 99 encoded an almost complete 16S rRNA gene sequence. A total of 23 gspp were identified as high-quality MAGs according to MIMAG standards (completeness ≥ 90% and contamination ≤ 5%^28^). The taxonomic classification based on GTDB and SILVA agreed, and both indicated that MAGs were affiliated with the classes *Gammaproteobacteria* (n=55), *Alphaproteobacteria* (n=14), *Acidimicrobiia* (n=17), *Acidobacteriota* class *Thermoanaerobaculia* (n=5) and *Planctomycetia* (n=3) (Table S8). The taxonomic classification of the recovered gspp revealed a significant level of novelty, as none of the 115 MAGs could be classified at the species level using GTDB. Only five MAGs could be assigned to known genera, including *Bythopirellula*, *Breoghania*, *Pseudocolwellia* and *Tateyamaria*, while 45 and 26 MAGs were affiliated with well-characterized families and orders, respectively (Table S7).

The combined relative abundance of all MAGs ranged from 7.4% to 11.4% of the total microbial community (in the Sep18 and Mar18_A4 samples, respectively) (Table S7). The observed abundances at class level were consistent with those detected using OTUs, with the *Gammaproteobacteria*, *Alphaproteobacteria*, and *Acidimicrobiia* classes being the most abundant (Figure S6). None of the MAGs independently reached an abundance > 0.82%, suggesting a lack of dominance by any single species (Table S7). Interestingly, the order *Woeseiales*, belonging to the *Gammaproteobacteria* class, was the most diverse (15 MAGs representing 8 genera and 2 families) and the most abundant in all metagenomes, averaging 1.3 ± 0.1%. Finally, the relative abundance of the only MAG belonging to the archaeum *Nitrososphaeria* was, on average, 0.3 ± 0.1%.

### Water column microbial diversity compared to sediment

To determine the similarities and differences between sediment and water column microbial communities, we examined the composition of OTUs determined from 16S rRNA sequences recovered from LR metagenomic datasets. Between March 19 and May 29 of 2018, we identified 993 and 621 OTUs in sediments and water column samples, respectively. When comparing all water column and sediment samples, we identified 276 OTUs shared between both fractions, but only nine OTUs were shared between all samples (Figure S7). During the same time frame but at the genomic level, 505 and 16 bacterial and archaeal representative gspp (based on MAGs) were recovered from water column metagenomes. These genomes were recovered from three size fractions (0.2-3, 3-10, 10+ µm), thus representing a proxy for free-living and particle-attached fractions (Table S8)^29^. A phylogenetic reconstruction using the 505 bacterial gspp from the sediment and water column revealed significant differences in microbial diversity between the two environments (Figure 2). While *Gammaproteobacteria* species were recovered from both sediment and water column metagenomes, the order *Woeseiales* was overrepresented in sediments and orders *Pseudomonadales* or SAR86 were predominant in the water column. Also, *Alphaproteobacteria* MAGs from sediments were predominantly assigned to *Rhizobiales*, while those from water column samples were related to *Pelagibacterales*. In water column metagenomes, the most diverse phylum was *Bacteroidota* (n=120), particularly within the *Flavobacteriales* order (n=101). In contrast, in sediments, only four MAGs were identified as *Bacteroidota*, two of which were affiliated with *Cytophagales*. *Verrucomicrobiota* and *Patescibacteria* were exclusively recovered from the water column.

**Figure 2.**
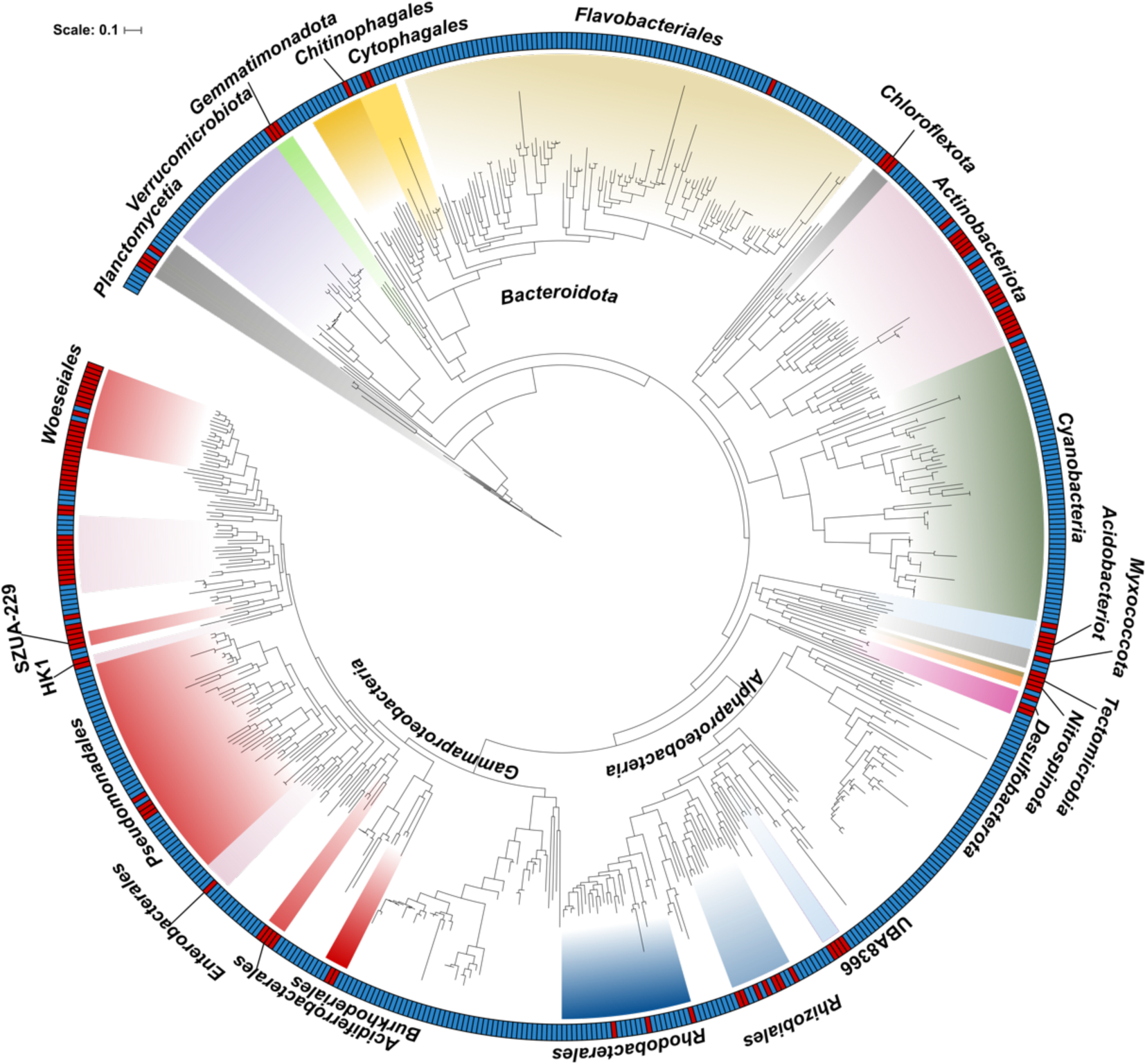
Phylogenetic reconstruction of the MAGs recovered from sediments and water column metagenomes in Helgoland. The maximum-likelihood tree and taxonomic classification were based on the conserved single-copy genes. Class and order names are indicated for MAGs recovered from sediment or water column metagenomic samples. Colored clades indicate the phylum, classes or orders indicated in the outer part of the phylogenetic tree. MAGs recovered from sediment (red) or water column (blue) samples are indicated in the external ring.

Only two MAGs from sediments were identified at the species level in the water column metagenomes. The Desulf_03 MAG shared a 99.2% ANI with the *Desulfocapsaceae* MAG GCA_905479735, and the Acti_15 shared a 98.1% ANI with the *Acidimicrobiia* MAG GCA_905480055. We mapped the short metagenomic reads from water column samples on the sediment-recovered MAGs to further corroborate their presence in water column metagenomic samples. The sediment Desulf_03 MAG had 28x sequencing depth (truncated average sequencing depth, TAD80) and a 99% sequencing breadth (i.e., the fraction of genome covered by metagenomic reads). The Acti_15 MAG only had ∼4.4x sequencing depth (TAD80) and 86% sequencing breadth. Interestingly, in the water column, both species were only identified in the particle-attached fractions metagenomes throughout the temporal series, with their maximum abundance observed in March and early April, coinciding with the pre- and initial stages of the spring algae bloom (Figure S8). Acti_15 was more abundant in the sediment metagenomes than Desulf_03, with an average relative abundance of 0.12 ± 0.04% and 0.03 ± 0.01%, respectively (Table S7).

### Organic matter degradation potential in water column vs sediments

To explore the potential contribution of sediment microbial communities in the degradation of organic matter, we compared the carbohydrate active enzyme (CAZymes) gene composition, including glycoside hydrolases (GH) as well as peptidases and sulfatases, among the representative MAGs from each gspp identified in both sediment and water column samples. On average, the fraction of genes encoding these polysaccharide degradation features was lower in sediment MAGs than those from the water column (Figure S9). Nonetheless, a higher fraction of GH and peptidase genes were encoded in both sediments and water column in *Bacteroidota* and *Gammaproteobacteria* MAGs, respectively. High sulfatase gene content was characteristic of *Planctomycetota* from the water column and *Bacteroidota* in sediments. While water column populations associated to the spring blooms are characterized for carrying polysaccharide utilization loci (PULs)^2,6,11,13,14^, only 17/115 gspp recovered from sediments carried PULs/CAZyme clusters (Table S9). In sediments, the most commonly detected PULs were associated with the degradation of laminarin (n=14), followed by alginate (n=9) and α-glucan (n=8). The majority of these PULs encoded in sediment MAGs were affiliated with the *Woeseiaceae* family (eleven MAGs; see below), *Bacteroidia* (three MAGs) and one *Acidobacteriota* (family *Thermoanaerobaculia*) MAG.

### Microbial diversity of Woeseiaceae family

The *Woeseiaceae* family presented the highest species diversity in the Helgoland sediments, represented by 50 OTUs at the 16S rRNA gene level and 14 MAGs at the genome level, and encoded the largest number of PUL-like structures identified in the recovered MAGs. We compared the phylogenetic context of the *Woeseiaceae* genomes recovered in the North Sea to those recovered from different sources (Figure 3). At first glance, we found that members of different genera were specific to their source. For instance, species of the genera UBA1847 clade I, JAACFE01, UBA1844, JAACFB01, IDS-8 and *Woeseia* were retrieved from sediments, coral reefs or marine biofilms (marked in red, pink and green in Figure 3, respectively). Contrastingly, species of genera UBA1847 clade II, SZUA-117, PBBK01, and SP4206 were predominantly recovered from water column samples (marked in blue in Figure 3). Distinctions in genome size between MAGs recovered from sediments and water column were observed (Figure S10), with sediment MAGs showing a larger genome size (average 3.1 ± 0.8 Mbp) compared to those from the water column (1.7 ± 0.7 Mbp).

**Figure 3.**
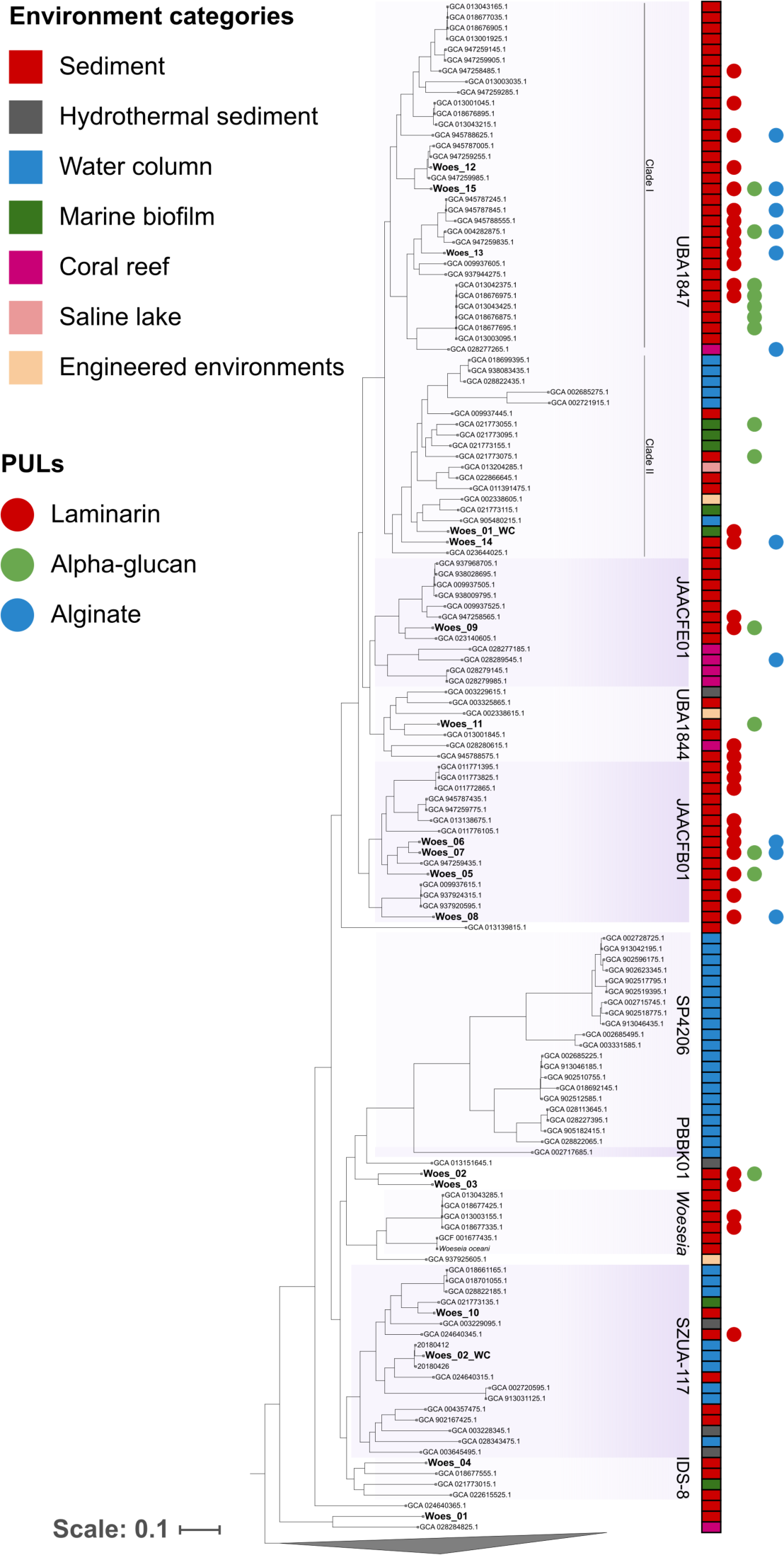
Phylogenetic reconstruction of the *Woeseiaceae* genomes using reference genomes, including MAGs recovered from sediment and water column metagenomes in Helgoland. Genomes marked in bold were retrieved from sediment and water column samples (this study), and others were obtained from NCBI and the Genome Taxonomy database. The colored squares indicate the source of each genome and MAG. The colored circles indicate the presence of laminarin (red dots), alpha-glucan (green), and alginate (blue) PULs.

To further understand the ecological role of *Woeseiaceae* species in sediments, we compared the MAGs recovered from sediments to those recovered from the water column at the North Sea^29^. In the Helgoland sediments, the Woes_10 (affiliated with SZUA-117) and Woes_08 (JAACFB01) were the most abundant *Woeseiaceae* species with average relative abundance of 0.28 ± 0.11% and 0.23 ± 0.1%, respectively (Figure S11). No *Woeseiaceae* species originating from sediments were detected in the overlaying water column metagenomes (detection based on read mapping, using a sequencing breadth cutoff of 20%). The *Woeseiaceae* MAGs Woes_01_WC (UBA1847 genus) and Woes_02_WC (SZA-117 genus), were identified in the particle-attached fraction but not in the free-living fraction or sediment samples (based on sequencing depth and breadth; Table S8 and Figure S12). These two water column MAGs had the highest abundance on March 19^th^ in the 3-10 µm, and in April 12^th^ in the 10+ µm fractions (Figure S12). Thus, when examining all samples recovered from the North Sea, we consistently observed a distinct environmental (i.e., niche) specificity of *Woeseiaceae* species to either sediment or water column samples, which is reflected in their metabolic potential.

### Metabolic potential of Woeseiaceae MAGs

#### Woeseiaceae heterotrophy through polysaccharide utilization loci (PUL) genes and peptidases

When examining *Woeseiaceae* genomes recovered from Helgoland and databases (NCBI and GTDB), *Woeseiaceae* genomes encoding PULs were identified only in sediment samples, as well as in particle-attached lifestyles such as marine biofilms and coral reefs (Figure 3). PULs were not identified in the two *Woeseiaceae* species retrieved from the water column samples (Woes_01_WC and Woes_02_WC). For instance, most of the genomes encoding PUL genomic contexts (e.g., targeting the degradation of laminarin, α-glucan and alginate) were clustered in the phylogenetic clade containing the genera UBA1847, JAACFE01, UBA1844, and JAACFB01. Moreover, *Woeseiaceae* MAGs obtained from sediment had the highest percentage of genes encoding for CAZymes and GHs when compared to MAGs recovered from water column (Figure 4).

**Figure 4.**
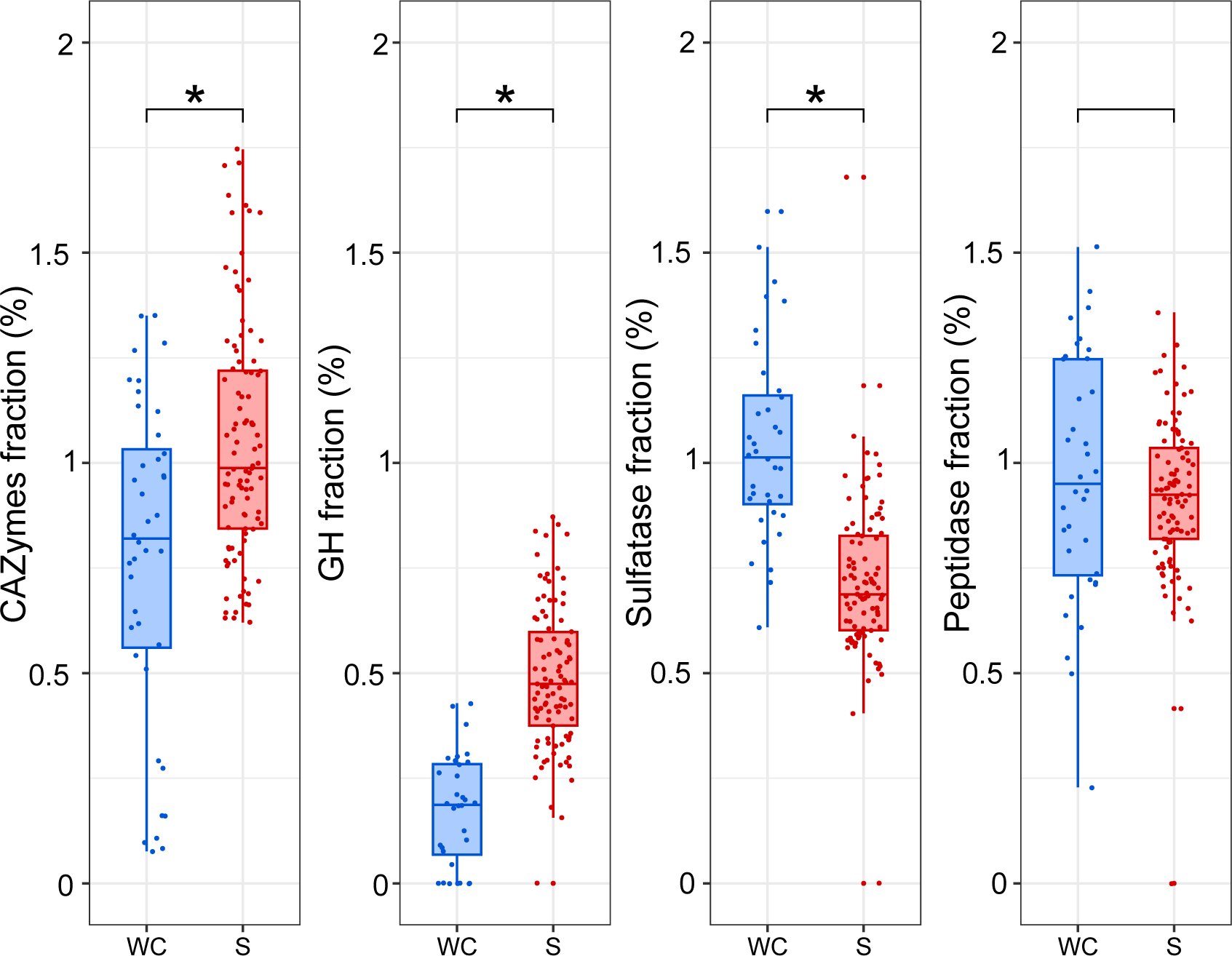
Fraction of genes encoding CAZymes, glycoside hydrolases (GH), sulfatases, and peptidases in *Woeseiaceae* MAGs according to their source. Box plots summarize the fraction of each *Woeseiaceae* MAG dedicated to different functional categories. Each point on the plot represents a single MAG and is colored blue or red according to a water column (WC) or sediment (S) origin, respectively. The asterisk denotes a statistically significant difference determined using the Wilcoxon test (p-value < 0.05).

We identified that ∼79% (11/14) of the recovered *Woeseiaceae* MAGs from Helgoland sediments encoded glycoside hydrolase (GH) genes associated with laminarin PULs, such as GH3, GH16, GH17, GH149, GH13, GH158 and GT51, with seven of these PULs containing a single starch utilization system SusC gene (Figure S13; Table S9). Six *Woeseiaceae* MAGs encoded genes for α-glucan PULs, characterized for the presence of GH13, GH31, GH97 and GH77 (Figure S14). Additionally, six MAGs encoded for alginate PULs, all of them carrying the PL6 and PL7 genes and including the PL17 or PL29 genes (Figure S15).

MAGs belonging to the *Gammaproteobacteria* class recovered from both water column and sediments encoded the highest fraction of peptidase-encoding genes, in particular, MAGs from sediments belonging to the *Woeseiales* and *Burkholderiales* orders and the *Pseudocolwellia* genus (*Enterobacterales* order) (Figure S9 and Table S10). Additionally, *Woeseiaceae* MAGs retrieved from the water column had a greater proportion of sulfatases and peptidases, supporting the idea that populations of the same family are specialized to different substrate niches (Figure 4).

#### Niche-specific Woeseiaceae metabolic patterns in different environmental systems

*Woeseiaceae* MAGs affiliated to SZUA-117 and UBA1847 clade II genera were retrieved from the water column and sediment in Helgoland metagenomes, and also from databases. Thus, we sought to examine the genetic components involved in niche-specific metabolic differentiation (Figure 5). Although some general metabolic pathways such as the TCA cycle and oxidative phosphorylation pathway were detected in all MAGs, source-specific metabolic pathways were also detected. For instance, the Entner-Doudoroff (ED) pathway was exclusively found in genomes from the UBA1847 genus recovered from sediments and marine biofilms. Similarly, genes associated with the electron transport complexes (ETC), such as the cytochrome c oxidase (cbb3-type) of the complex IV were encoded only in MAGs from sediment samples or marine biofilms. Incomplete denitrification pathways were detected in most genomes from sediments, carrying the genetic potential for the nitrate reductase NapAB (reduction of nitrate to nitrite), nitrite reductase NirS/NirK (reduction of nitrite to nitric oxide) and/or the nitric oxide reductase NorBC (reduction of nitric oxide to nitrous oxide). Rubisco (ribulose-bisphosphate carboxylase large chain) required for carbon fixation via the pentose phosphate cycle, was identified only in the sediment MAG Woes_15 suggesting autotrophic metabolism for the minority of the *Woeseiaceae* MAGs recovered (Table S12). Interestingly, the dissimilatory sulfate reduction or the conversion of thiosulfate to sulfite was only detected in sediment and marine biofilm genomes. In terms of phospholipid membrane transport, we observed that the nearly complete Mla (maintenance of outer membrane lipid asymmetry) pathway was predominantly present in genomes from sediment samples. Furthermore, most sediment MAGs exhibited a complete set of genes associated with flagellar assembly. The twitching motility genes (PilTIJU) were encoded in all Helgoland *Woeseiaceae* MAGs recovered from sediments and water column. The gene encoding for the tight adherence (TadA) pilus was identified in three MAGs retrieved from Helgoland sediments (Woes_01, Woes_07, and Woes_11), while it was absent in the *Woeseiaceae* MAGs obtained from the water column (Table S12). Overall, the comparative analysis of functional genes among *Woeseiaceae* MAGs retrieved from the water column and sediment indicated a better adaptation of sediment *Woeseiaceae* species to low-oxygen or anoxic conditions.

**Figure 5.**
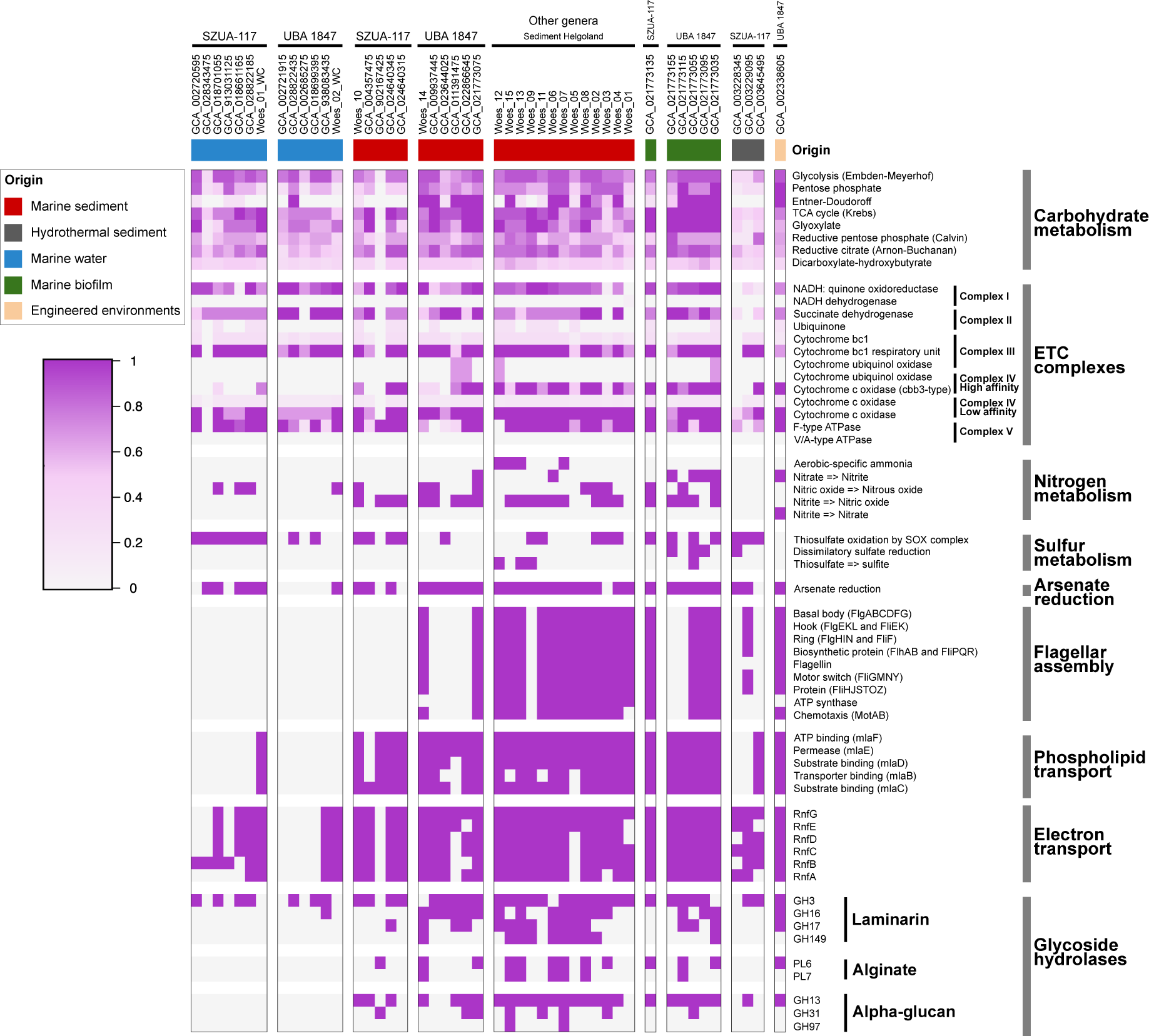
Summary of the metabolic potential for *Woeseiaceae* genomes and MAGs affiliated to genus SZUA-117 and UBA1847. The genomes and MAGs are organized based on their origin.

## Discussion

### LRs sequencing considerations in sediment samples

The examination of the metagenomic data recovered here suggested that replicates sequenced from the same DNA and a different DNA extraction of the same sample had the lowest dissimilarity values (based on MASH distance and taxonomic profiles) compared to samples originating from different seasons. The results confirmed the high seasonal stability of coastal marine sediment microbiomes. Moreover, the LR metagenomic approach allowed the retrieval of a large collection of high-quality and full-length 16S rRNA sequences from the benthic microbial community, which will facilitate further studies interested in designing FISH probes for microscopy-based identification. Despite the high diversity of benthic microbiomes, we could retrieve high-quality MAGs using long-read sequencing and co-assembly. Thus, our approach benefited from a high sequencing effort, the temporal stability of microbial communities, and the co-assembly of metagenomic samples for studying complex microbial diversity^30^. Nonetheless, assembling multiple samples might conceal intra-species diversity changes and only represent a smaller fraction of the metagenomic samples. The recovered MAGs represented approximately 7-11% of the total microbial community, primarily due to the lower sequencing effort per run in LR sequencing that results in less coverage of the microbial diversity^27^. Nonetheless, the LR metagenomic approach represents a leap forward for recovering medium to high-quality MAGs from complex environmental samples.

### Temporal stability of sediment microbial communities and interaction with overlaying water column samples

We found a clear phylogenetic separation of MAGs recovered from the sediment and the water column. Together with the presence of specific metabolic pathways, our findings highlight an evolutionary imprint reflected in the genetic specializations and adaptations for bacterial populations recovered from sediments and the overlaying water column (Figure 2). At the species level, only a small fraction of populations overlapped in both the water column and sediment metagenomes (e.g., *Acidimicrobiia* Acti_15 and *Desulfocapsaceae* Desulf_03). We hypothesize that both species originated from the sediment and were subsequently upwelled into the water column, as they were exclusively identified in the particle-attached fractions (3-10 and 10+ µm) and not in the free-living fractions (0.2-3µm). During spring, the water column is generally higher in turbidity compared to the bloom phase^31^, suggesting the resuspension of sediments, and thus explaining the detection of sediment microbial species in the particle-attached water column samples. Moreover, based on the OTU analysis, we identified the Archaea *C.* Nitrosopumilus as the most dominant single species in sediments, likely missed in previous studies relying only on Bacteria domain-specific primers^9^. In fact, members of the phylum *Nitrososphaerota* (formerly Thaumarchaeota) have been identified in oxic sediments in various marine environments, including shallow estuaries, open oceans and deep oceanic crust^32,33,34,35^. The microbial composition and temporal stability of benthic microbial assemblages in Helgoland sediment assessed here through deeply sequenced LRs metagenomes, matches the findings derived from amplicon 16S rRNA gene sequencing and microscopy cell counts from the same samples^9^. This stability in benthic bacterial communities has been observed in other ecosystems, such as those in Svalbard (Arctic Ocean^9^) or at the island of Sylt (North Sea^36^). While the stability of species abundance is evident in sediments, particularly within the *Woeseiaceae* family, it is noteworthy that these organisms might exist in a latent state for an extended period of time, awaiting opportune conditions (both biotic and/or abiotic) to stimulate their metabolic activity and exploit their ecological niches.

### Metabolic profiling and niche specialization of Woeseiaceae species in sediments and water column ecosystems

*Woeseiaceae* family members have been recognized as highly diverse and abundant microorganisms across various sediment environments, including oxic surface coastal sediments, sublittoral, hydrothermal vent, and deep-sea locations^9,15,17,21,23,25,37^. In this study, we observed that all *Woeseiaceae* species were retrieved exclusively from marine environments, including pelagic environments, particle-attached fractions, or from marine benthos, but not from freshwater environments. Despite their prevalence in marine sediments, their diversity and metabolic potential remain largely understudied. The metabolic analysis of the *Woeseiaceae* MAGs recovered here supports the chemolithoautotrophy potential based on hydrogen (presence of oxygen-tolerant [NiFe]-hydrogenase) and sulfur for thiosulfate oxidation (encoding SOX gene system) suggested before^21,23,24,25^. Moreover, the presence of a Rubisco large chain in Woes_15 MAG is indicative of carbon fixation using the Calvin Benson cycle, which was also identified in MAG JSS_woes1 recovered from Janssand tidal sediment^25^.

Members of *Woeseiaceae* seems to be mixotrophs. Their heterotrophy is linked to the remineralization of polysaccharides and proteins. Here, we found numerous CAZyme genes organized in PULs and peptidases encoding genes^38^. Peptidase genes were enriched in *Woeseiaceae* MAGs from the water column, whereas sediment *Woeseiaceae* MAGs encoded a higher proportion of GH genes (Figure 4). For example, *Woeseia oceani* has been previously defined through the large diversity of its peptidases, emphasizing their proteolytic potential^15,21,25^. Our long-read sequencing approach allowed us to recover a PUL organization of CAZyme genes, transporters, and regulators often lost in highly fragmented MAGs recovered using short-read sediment metagenomes. Unlike the PULs frequently found in *Bacteroidota*^30^, the PUL structures in *Gammaproteobacteria* were distinguished by the absence of SusCD genes tandem^40^. The lack of typical PUL structures has also been observed in *Verrucomicrobiota* MAGs lacking SusCD genes^13^. Then, the sediment *Woeseiaceae* species encoded for PUL structures, which were not encoded in the water column *Woeseiaceae* MAGs, revealing their niche specialization for polysaccharide breakdown, and distinguishing them from the water column species that feature more sulfatase and peptidase genes.

The phylogenetic reconstruction indicated a differential representation of *Woeseiaceae* genera and species across environmental categories (Figure 3). A significant fraction of *Woeseiaceae* species were derived from sediments or other particle-attached environments. Nevertheless, species associated with pelagic environments were only affiliated to the genera UBA1847 clade II, SZUA-117 and SP4206, indicating an ecological niche specialization. This strong correlation between phylogenetic *Woeseiales* lineages and their source was previously observed by a phylogenetic analysis based on 16S rRNA genes retrieved from the SILVA 128 database^21^.

In the sediment and water column samples from Helgoland, we retrieved MAGs representing *Woeseiaceae* species affiliated with the UBA1847 clade II and SZUA-117 genera, and an evident metabolic divergence was observed depending on their source (Figure 5). Most MAGs recovered from sediments encoded a complete or almost complete ED pathway, which was not detected in water-column MAGs. The ED pathway provides sediment *Woeseiaceae* species with a versatile metabolism for using glucose under oxic and/or anoxic conditions. Compared to Embden-Meyerhof-Parnas pathway, the ED requires fewer enzymatic steps, enhancing metabolic efficiency and avoid the production of reactive oxygen species (ROS) under anoxic conditions^41,42^. Moreover, the sediment MAGs encoded the high affinity complex IV of the ETC, including the cytochrome c oxidase cbb3-type, indicating their potential to respire under low-oxygen or anoxic conditions, and also to perform nitrate/nitrite respiration, depending on prevailing environmental conditions^43^. Under anoxic or low-oxygen conditions, gram-negative bacteria such as *Woeseiaceae* may experience stress that could disrupt the lipid asymmetry of the outer membrane. We observed that the MAGs recovered from sediments encoded the Mla system that helps to restore and maintain this asymmetry by removing excess phospholipids from the outer membrane and transporting them back to the inner membrane^44^. In contrast to water column *Woeseiaceae* MAGs, the presence of flagella and the tight adherence (tad) pilus genes in sediment-dwelling species may facilitate movement to favourable environments, promote surface adhesion, and enhance colonization and biofilm formation^45,46,47^. Moreover, all *Woeseiaceae* MAGs, whether from pelagic or benthic environments, encoded for twitching motility genes (type IV pili) that facilitate cell-cell and cell-surface adhesion^48^.

## Methods

### Study site and sampling

Samples were obtained from a shallow coastal site at Helgoland Roads, North Sea (German Bight; 54.18°N, 7.90°E), with water depths ranging over the tidal cycle between ∼3 and ∼8 m. Sediment sample collection and description were previously detailed^9^. Briefly, sediment samples were obtained monthly by scientific divers from the Alfred Wegner Institute (Bremerhaven, Germany) using push cores. In this study, we selected five samples collected on March 7th 2018 (referring as sample Mar18), April 17th 2018 (Apr18), May 15th 2018 (May18), September 8th 2018 (Sep18) and January 18th 2019 (Jan19). DNA was extracted from the top 2 cm of the sediment surface as reported by Miksch et al. (2021)^9^.

### Metagenome analyses using PacBio long-reads

The extracted DNA was quantified using fluorometry (Quantus, Promega), and the quality was assessed using capillary electrophoresis (Agilent FEMTOpulse). Metagenomic libraries were prepared using the HiFi SMRTbell® Libraries from Ultra-Low DNA Input protocol. HiFi SMRT sequencing was done for 30 hours on a Sequel IIe device at the Max Planck Genome Center in Cologne, Germany. Metagenomic LR quality was assessed using Nanoplot v1.39.0^49^. Beta-diversity analysis was conducted using MASH distance (Mash v1.1^50^. MASH distance values were visualized in an NMDS plot using the vegan library^51^. Taxonomic classification of metagenomic LRs was performed using Kaiju v1.6.3^52^.

16S rRNA genes were extracted from metagenomic LRs using barrnap v0.9 (https://github.com/tseemann/barrnap) and clustered in operational taxonomic units (OTUs) at 98.7% similarity (minimum threshold discriminating species suggested by Stackebrandt and Goebel (1994)^53^) using uclust v1.2.22^54,55^. Singleton and doubleton OTUs were removed from the analysis. Representative OTU sequences were imported into the non-redundant SILVA REF 138 database^56^. Sequences were aligned using the SINA v1.3.1 aligner^57^ implemented in the ARB program package v6.0.6^58^. Rarefaction analyses and alpha diversity indices were conducted using PAST v4.03^59^.

### Metagenomic assembly and binning

To compare the effect of sequencing effort with the number of recovered of MAGs in sediment samples, we employed two co-assembly strategies that relied on the subsampling of the metagenomic dataset generated in this study (n=9). The first approach involved concatenating the metagenomic files in chronological order, while the second approach entailed concatenating all samples and subsampling to match the Gbps size of the previous strategy. Single and concatenated metagenomes were assembled with Flye v2.8.1^60^ using the options *pacbio-hifi* and *meta*. For each metagenomic dataset, contig sequencing depth (or coverage) was calculated mapping long-reads using minimap2 v2.17^61^ with the *map-hifi* option and using the *jgi_summarize_bam_contig_depths* script from MetaBAT v2.2.15^62^. Contigs larger than 2,000 bps were binned using MaxBin v2.2.7^63^ and MetaBAT v2.2.15^62^. MAG completeness and contamination values were calculated using CheckM v1.1.6^64^ and CheckM2 v0.1.3^65^. The quality for each MAG was estimated as the composite index ‘completeness – 5 x contamination’ and MAGs with quality ≥ 50 were selected for further analysis. MAGs obtained from the co-assembly of all samples generated in this study (n=9) were used for comparative analyses. dRep v3.2.2 was used to de-replicate MAGs in gspp with the options -pa 0.96 (ANI threshold to form primary clusters)^66^. Gene prediction from LR metagenomes was conducted using Prodigal v2.6.3^67^, and representative gspp MAGs were selected for abundance estimation.

### MAGs abundance estimation using PacBio long reads

To measure the average number of universally conserved single-copy genes in the LRs metagenomes (i.e., genome equivalents), we used the script *HMM.essential.rb* (with the “metagenome” option) from enveomics collection^68^. Genome equivalents were calculated for each metagenomic dataset as the average number of genes predicted for each set of universally conserved single-copy genes. The sequencing depth for each MAG was estimated mapping the unassembled LRs using minimap2 v2.17^61^ with the *map-hifi* option, and filtering mapped LRs at ≥ 95% identity using the *jgi_summarize_bam_contig_depths* script from MetaBAT v2.2.15^62^. These average sequencing depth values were normalized using the genome equivalents determined for each metagenome^27^.

### Metagenomes and MAGs recovery from water column metagenomes

We used metagenomes and MAGs from the overlaying water column samples recovered during the spring bloom in 2018^29^. All MAGs were dereplicated using dRep v3.2.2^66^ with a 96% ANI cut-off. To estimate MAG relative abundances, metagenomic reads were mapped against MAGs using the BLASTn option of the aligner Magic-BLAST+ v2.12.0 with default settings^69^. Reads matching with higher than the 95% sequence identity and an (alignment length) / (query read length) greater than 0.9 were used to determine the sequencing depth for all representative MAGs in each metagenomic dataset. The resulting read depth values were truncated to the middle 80% (TAD80) of depth values (i.e., the upper and lower 10% of outliers were removed) using a Python script from https://github.com/rotheconrad/00_in-situ_GeneCoverage ^13,70^. Finally, to estimate the relative abundance with respect to the abundance of bacterial and archaeal communities, each TAD80 value was normalized by the “genome equivalents” value estimated using MicrobeCensus v1.1.0^27,71^.

### MAGs phylogenetic classification

Phylogenetic analyses of MAGs were performed using the GTDB-tk v2.1.1 (release 207_v2) with the *classify_wf* pipeline^72^. The alignment of housekeeping proteins produced in GTDB-tk was used for phylogenetic tree reconstruction using IQ-TREE v1.6.12^73^. Visualization and editing of the phylogenetic trees were conducted using iTol^74^.

All 16S rRNA genes were extracted from LR MAGs using barrnap v0.9 (https://github.com/tseemann/barrnap). The sequences were imported into the latest updated Living Tree Project database (LTP_06_2022) containing all sequences of the type strains classified reported until June 2022^75^ and in the SILVA REF 138.1 database^56^. Sequences were aligned using SINA v1.3.1^57^ in ARB v6.0.6^58^, and manually checked to improve the alignment and to finally perform the phylogenetic analyses using ARB package v6.0.6^58^.

The ANI and fraction of genome shared between all MAGs was calculated using *ani.rb*^68^ based on the BLASTn^76^.

### MAGs gene annotation and PUL prediction

Protein-coding genes for all MAGs were annotated using the SwissProt and TrEMBL databases (downloaded in September 2022^77^) and DIAMOND v0.9.31 with default settings^78^. Also, proteins were annotated using the Kyoto Encyclopedia of Genes and Genomes (KEGG) database^79^ and DRAM v1.0 (Distilled and Refined Annotation of Metabolism)^80^. Annotations of CAZymes were done using the dbCAN v10 database^81^ and DIAMOND searches^78^ against the CAZy database v04242021 (E-value ≤ 1e-20)^82^. Only genes positive against both dbCAN and the CAZY database were considered reliably annotated as CAZymes. SulfAtlas v2.3.1^83^ was used to annotate sulfatases using DIAMOND (E-value ≤1e-4). The MEROPS v12.4 database^84^ was used to annotate peptidases using DIAMOND (E-value ≤1e-4). TonB-dependent transporters (TBDTs) were predicted by HMMscan against TIGRFAM profiles TIGR01352, TIGR01776, TIGR01778, TIGR01779, TIGR01782, TIGR01783, TIGR01785, TIGR01786, TIGR02796, TIGR02797, TIGR02803, TIGR02804, TIGR02805, TIGR04056 and TIGR04057 (E-value ≤1E−10). SusD genes were identified by HMMscan against the Pfam profiles PF12741, PF12771, PF14322, PF07980.

### Data availability

The metagenomes and MAGs generated in this study were deposited under the European Nucleotide Archive (ENA) accession code PRJEB64856, and accession run codes from ERR11809028 to ERR11809036 (Table S1). The metagenomes and MAGs from the three water column fractions are available under the accession code PRJEB38290^29^. The 16S rRNA sequences recovered from PacBio raw reads dataset are available in doi:10.6084/m9.figshare.25577322.

## Acknowledgements

The Max Planck Society funded this study. We acknowledge Christoph Walcher, Markus Brand, Madlen Friedrich, and the team for sampling at Helgoland (Centre for Scientific Diving), Antje Wichels, and Eva-Maria Brodte for providing infrastructure at Alfred Wegener Institute Helgoland. Metagenomic sequencing was done at Max Planck Genome Center in Cologne (Germany), supervised by Bruno Huettel. We thank Isabella Wilkie for her helpful suggestions regarding the manuscript. Tomeu Viver acknowledges the “Margarita Salas” postdoctoral grant, funded by the Spanish Ministry of Universities, within the framework of Recovery, Transformation and Resilience Plan, and funded by the European Union (NextGenerationEU), with the participation of the University of Balearic Islands (UIB).

## Author Contributions Statement

L.H.O. and R.A. designed and supervised the project. L.H.O. and T.V. conceptualized the project. T.V. performed all data analysis. T.V. and L.H.O wrote the manuscript. K.K. and R.A. reviewed and edited the manuscript. All authors read and approved the manuscript.

