## Supplemental Figures for "Deep long-read metagenomic sequencing reveals niche differentiation in carbon cycling potential between benthic and planktonic microbial populations"

**Affiliations**

**Figure S1. Relatedness between all metagenomic samples recovered.** The Non-metric Multidimensional Scaling (NMDS) analysis is based on MASH-based distances calculated between all pairs of metagenomic reads.

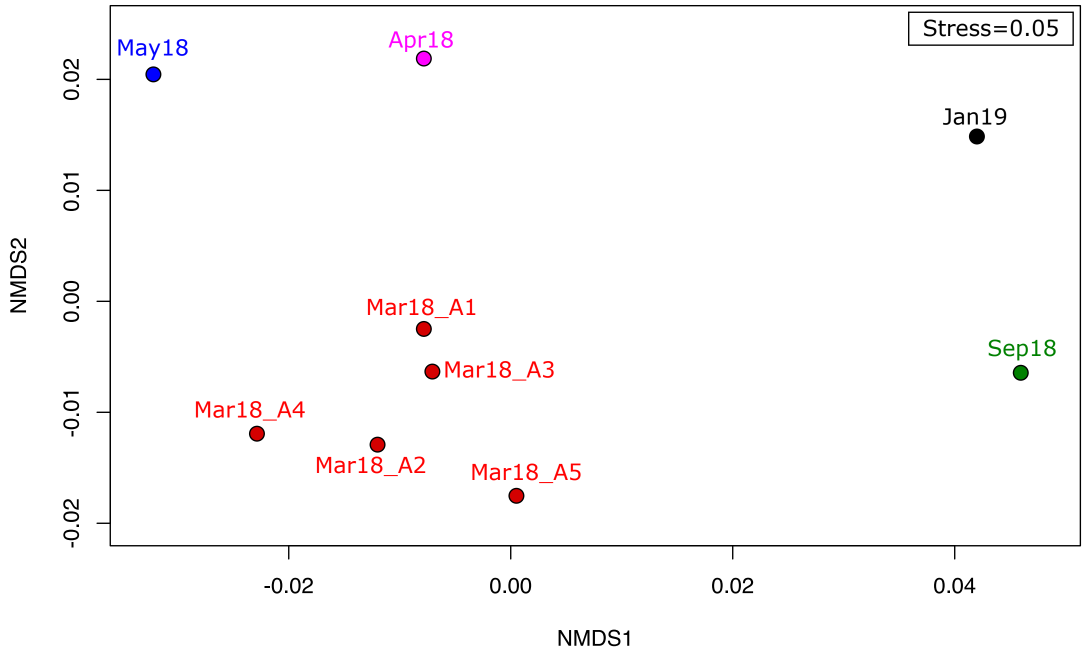

**Figure S2. Rarefaction curves based on Bacteria and Archaea OTUs classification of all metagenomic samples.** The x-axis denotes the number of 16S rRNA sequences recovered from each metagenomic dataset, while the y-axis corresponds to the number of OTUs (Richness). (A) Rarefaction curve for each individual metagenome. (B) Rarefaction curves for combined datasets: one for the five Mar18 samples (Mar18_Combined_dataset) and one for all metagenomes combined (ALL_Combined_dataset). Singleton and doubleton OTUs were removed from the analysis.

A

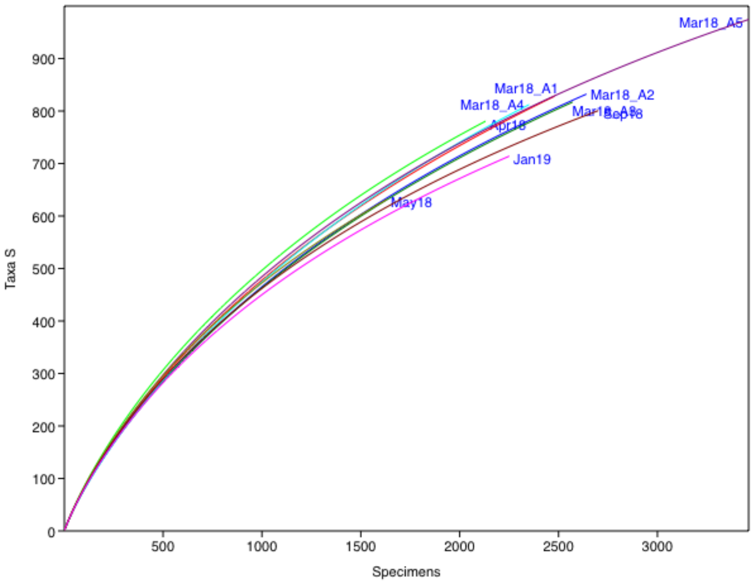


B

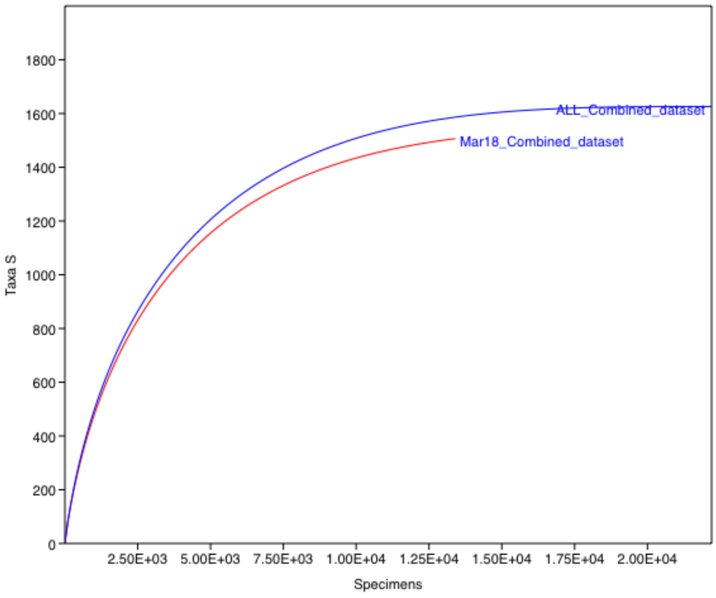

**Figure S3. Taxonomic distribution and relative abundance of bacterial and archaeal families and genera from Helgoland sediments derived from recovered 16S rRNA sequences from metagenomes.** Taxonomic classification is based on the SILVA SSU138.1 NR99 database. The circle diameter indicates the relative abundance (16S rRNA read frequencies) and color of the sampling season.

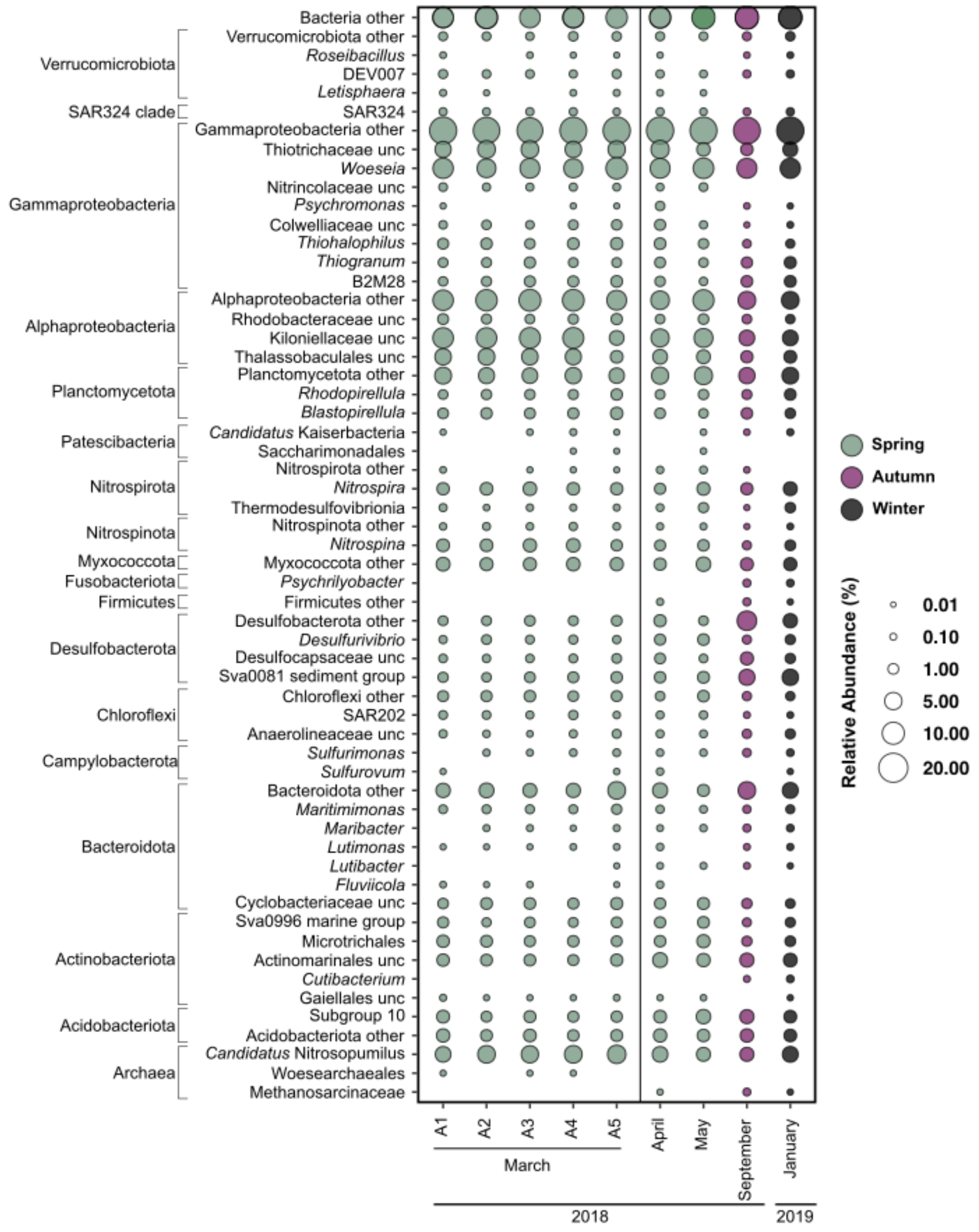

**Figure S4.** Comparison of **relative abundance values for the most abundant class and order levels in Helgoland sediments.** The results presented here compare the abundances determined using**:** (i) full-length 16S rRNA gene sequences from unassembled PacBio long reads and taxonomic classification based on OTUs, (ii) the taxonomic classification of unassembled metagenomic data reads based on sequence similarity (Kaiju) and (iii) 16S rRNA amplification approach reported in Miksch et al., 2021.

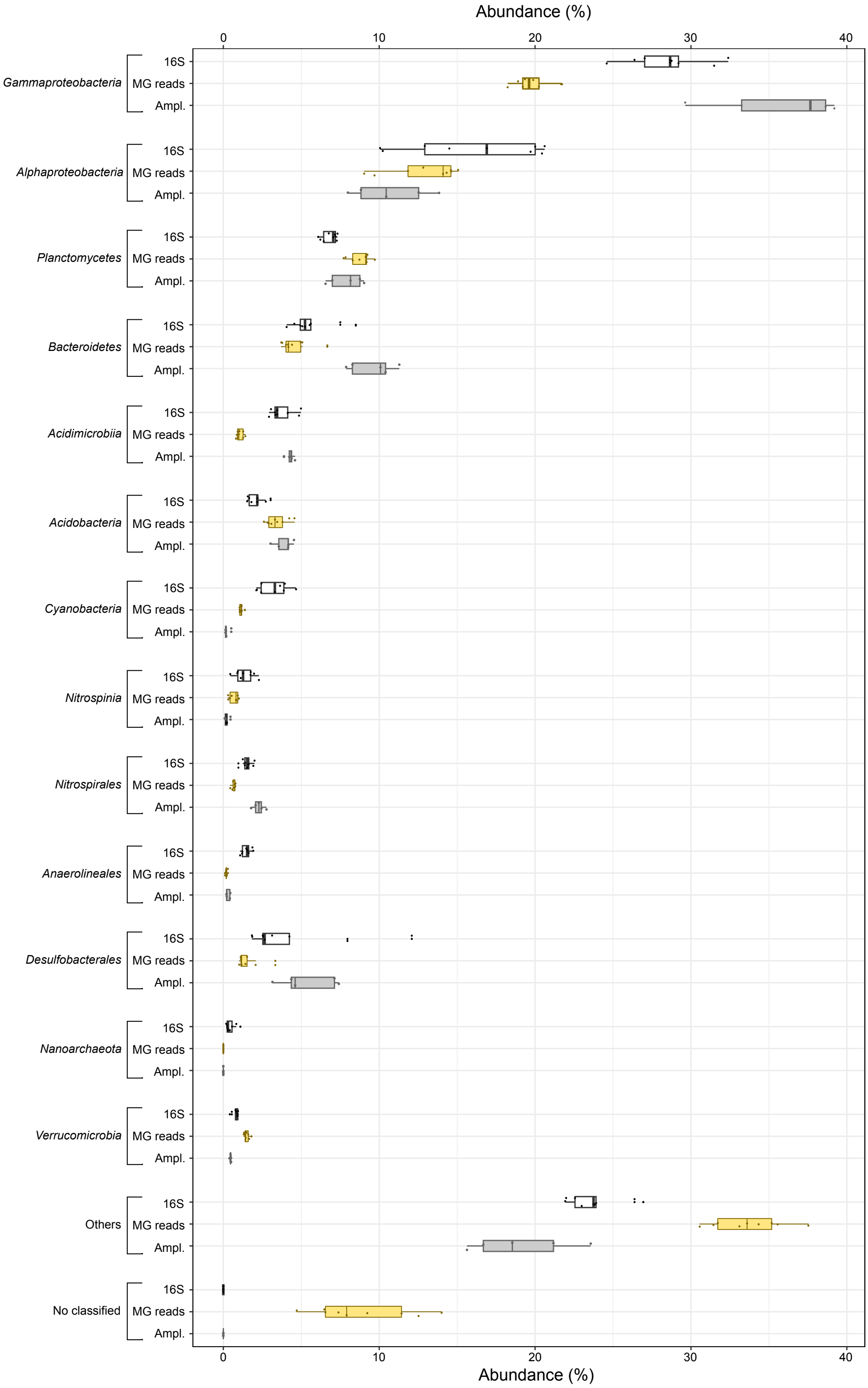

**Figure S5. Comparison of relative abundances determined using metagenomic and amplicon sequencing for sediment samples.** Comparison of the taxonomic distribution and relative abundances of bacterial and archaeal families and genera reported in Figure S3 using 16S rRNA gene sequences extracted from unassembled LRs and from amplicon sequencing previously reported (Miksch et al., 2021). Statistical significance was determined using the Wilcoxon test (p-value <0.05).

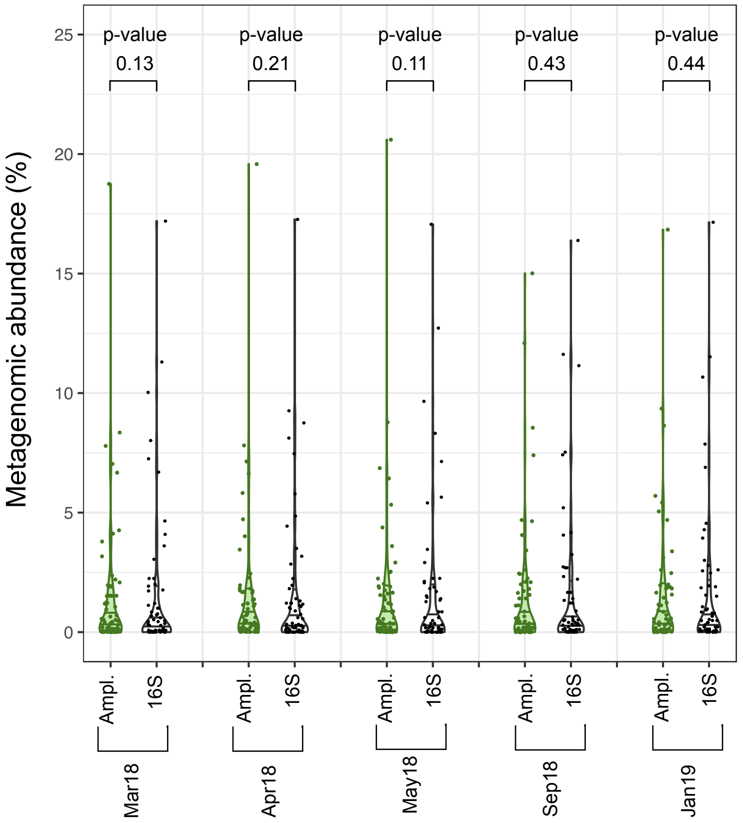

**Figure S6.** Relative abundance of MAGs, grouped at class level, retrieved from sediment samples. Note that from sample Mar18, five replicates from the sample were sequenced (A1 to A5).

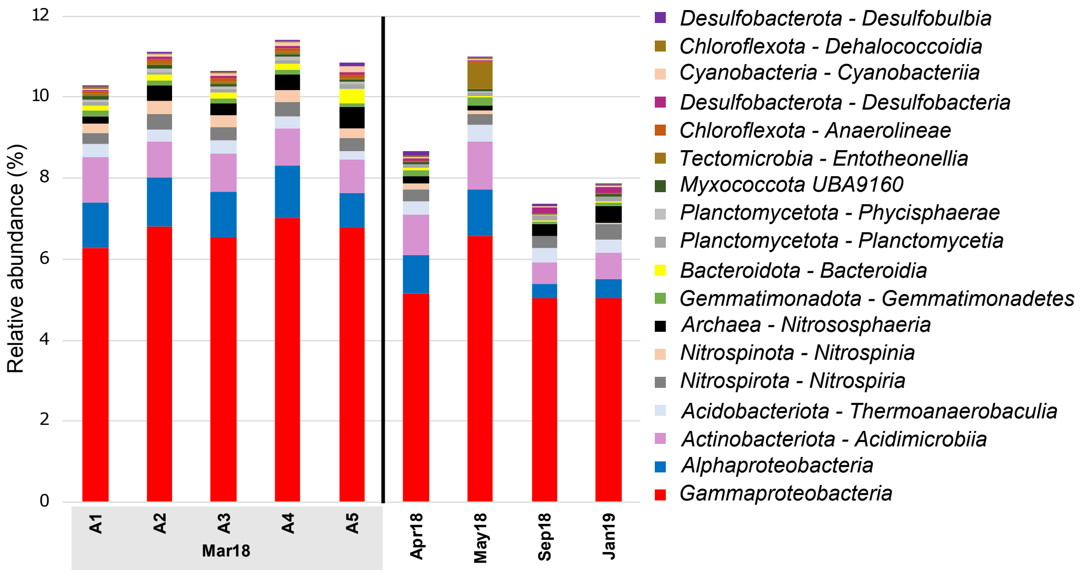

**Figure S7. Comparison of OTUs recovered from sediment and water column metagenomic samples collected between March 19 and May 29 of 2018.**

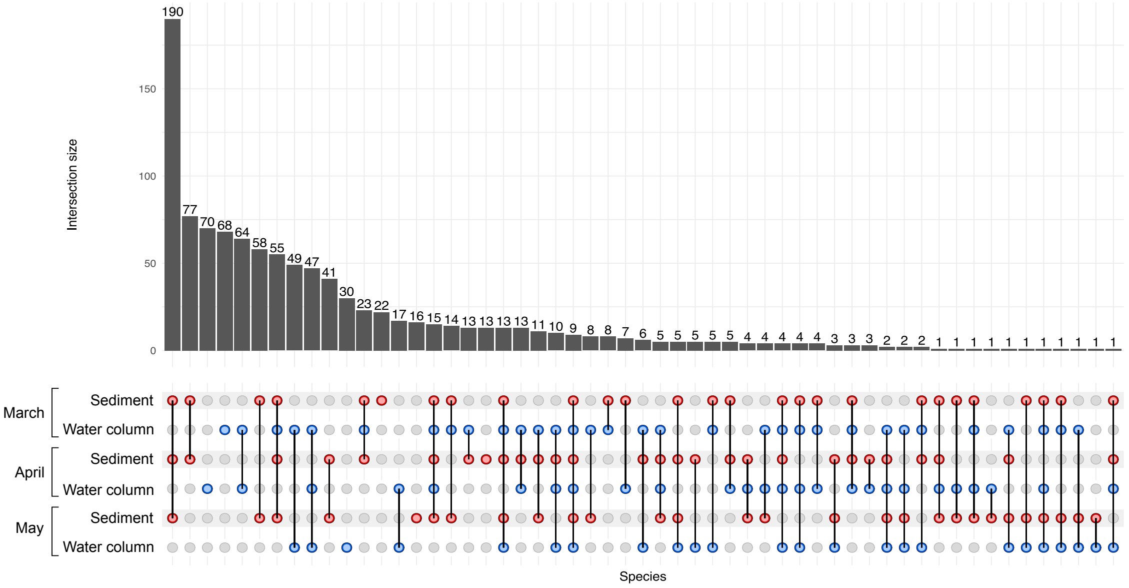

**Figure S8.** Relative abundance of the Desulf_03 and Acti_15 MAGs, affiliated to the family *Desulfocapsaceae* and class *Acidimicrobiia*, respectively. Their relative abundance was analyzed in the three distinct filtered fractions of the water column samples in 2018 (0.2–3 μm, 3–10 μm and >10 μm fractions) during three months. In samples from 2018-03-19 and 2018-04-12 and the 3–10 μm and >10 μm fractions, the *Desulfocapsaceae* MAG was identified with a sequencing breath (i.e., the fraction of the genome covered by metagenomic reads) >0.96. The *Acidimicrobiia* MAG had a sequencing breath >0.84 in the same samples.

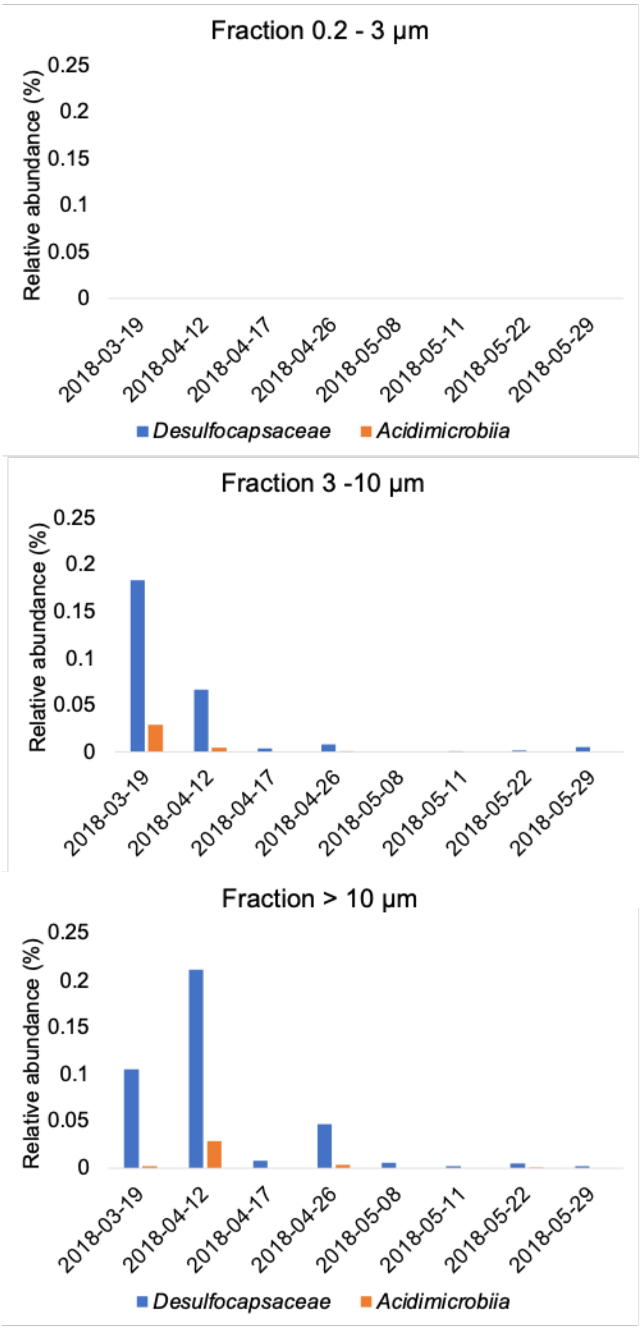

**Figure S9.** Fraction of genes encoding CAZymes, Glycoside hydrolases (GH), sulfatases, and peptidases in MAGs retrieved from the water column (blue) and sediment samples (red). Every point on the graph represents a single MAG, classified at the class level based on the phylogenetic reconstruction using the marker genes implemented in GTDB-tk tool.

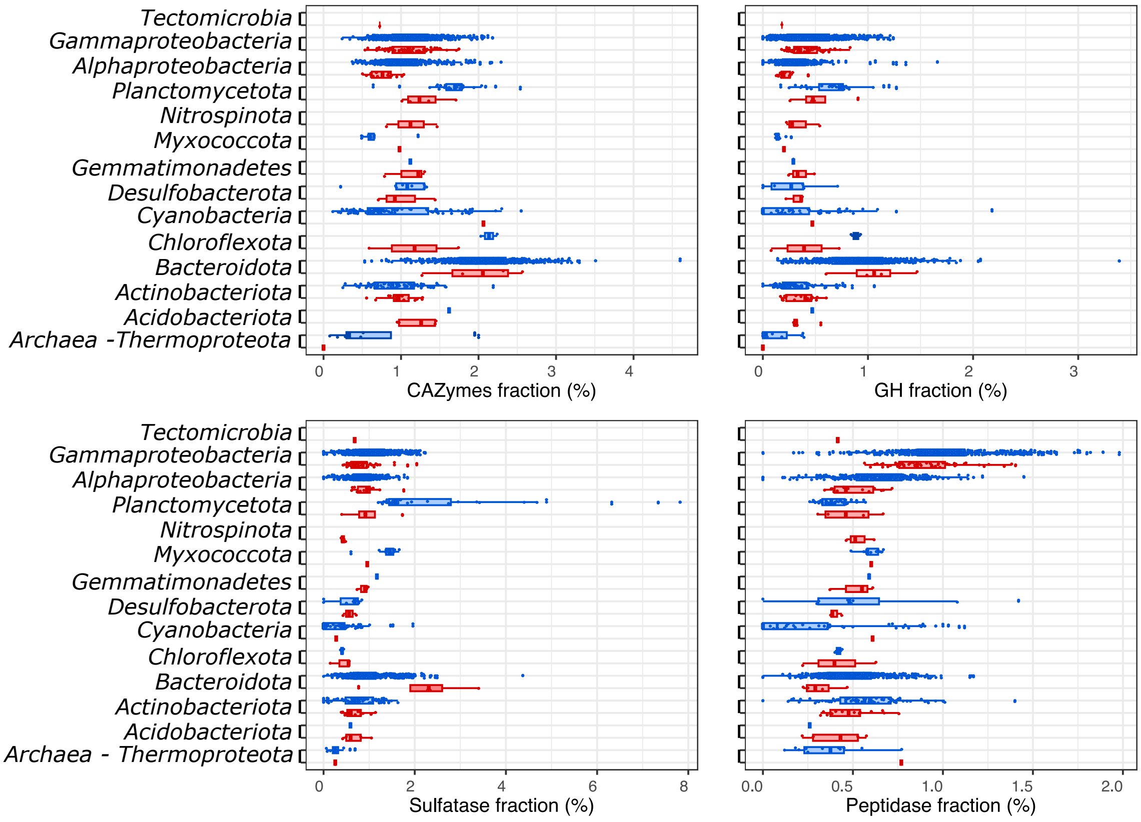

**Figure S10. Genome size of the *Woeseiaceae* MAGs classified depending on their water column (WC) or sediment (S) origin.** Every point on the graph represents a single MAG and is color-coded based on the environment categories from which it was retrieved. The asterisk denotes a statistically significant difference determined by the Wilcoxon test (p-value < 0.05).

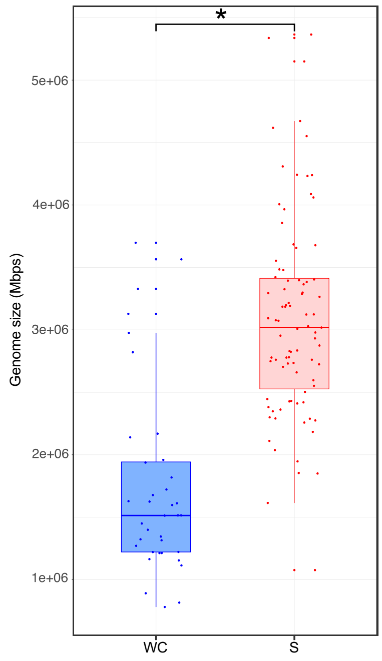

**Figure S11. Relative abundance of *Woeseia* MAGs in the sediment samples.** The y-axis represents the average abundance of each MAG in the temporal series in sediment metagenomes and its standard deviation.

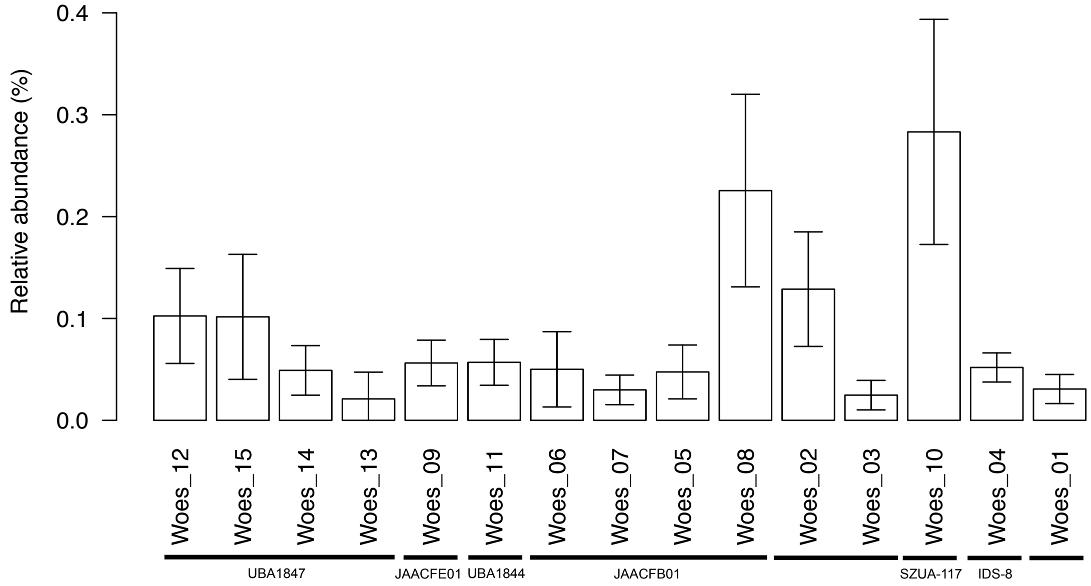

**Figure S12: Relative abundance of the *Woeseiaceae* MAGs retrieved from water column metagenomes**. Their relative abundance was analyzed in the three distinct filtered fractions of the water column samples in 2018 (fractions 0.2–3 µm, 3–10 µm and >10 µm) and in the sediment samples.

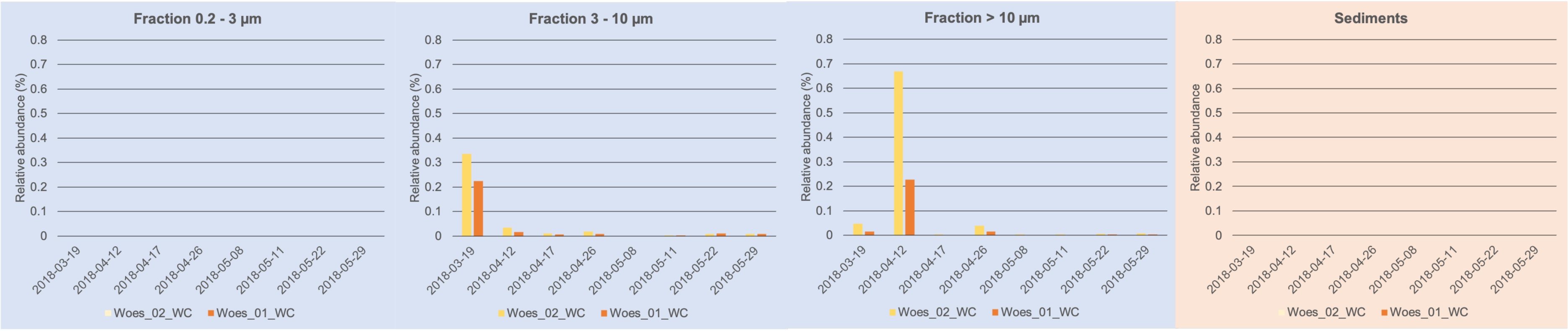

#### Figure S13: Genetic organization of the laminarin PULs encoded by *Woeseiaceae* MAGs.

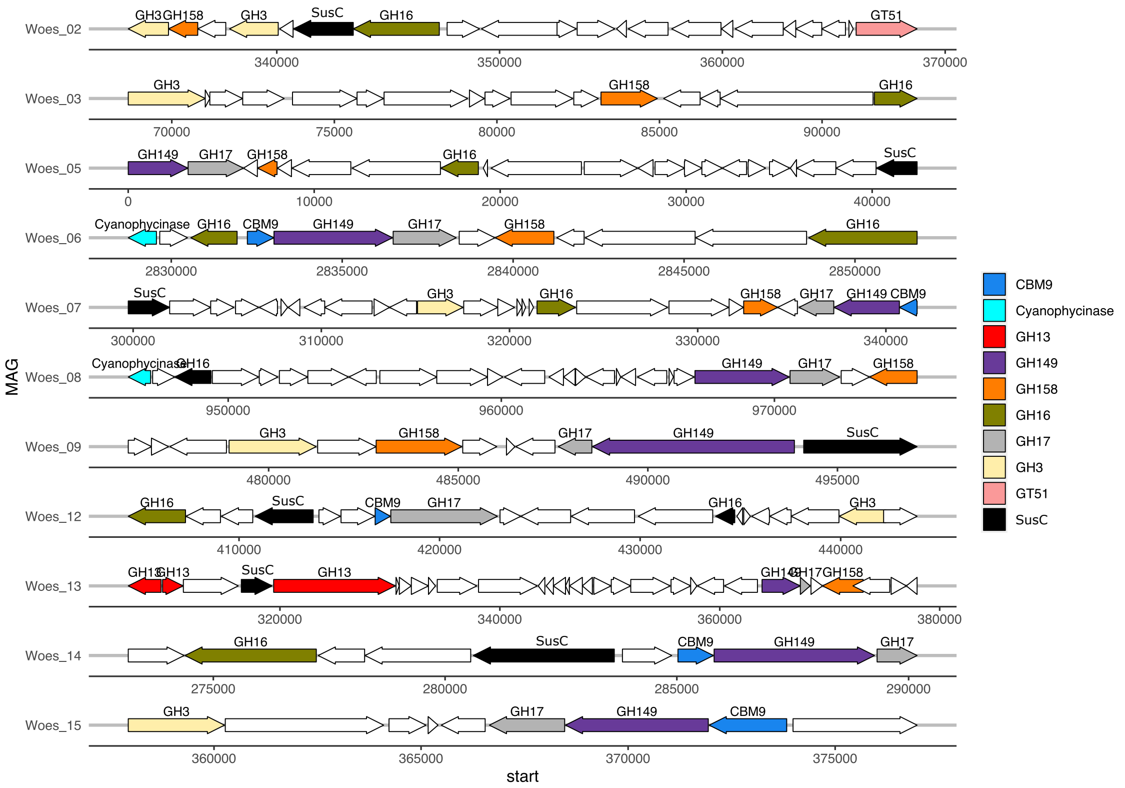

### Figure S14: Genetic organization of the alpha-glucan PULs encoded by *Woeseiaceae* MAGs.

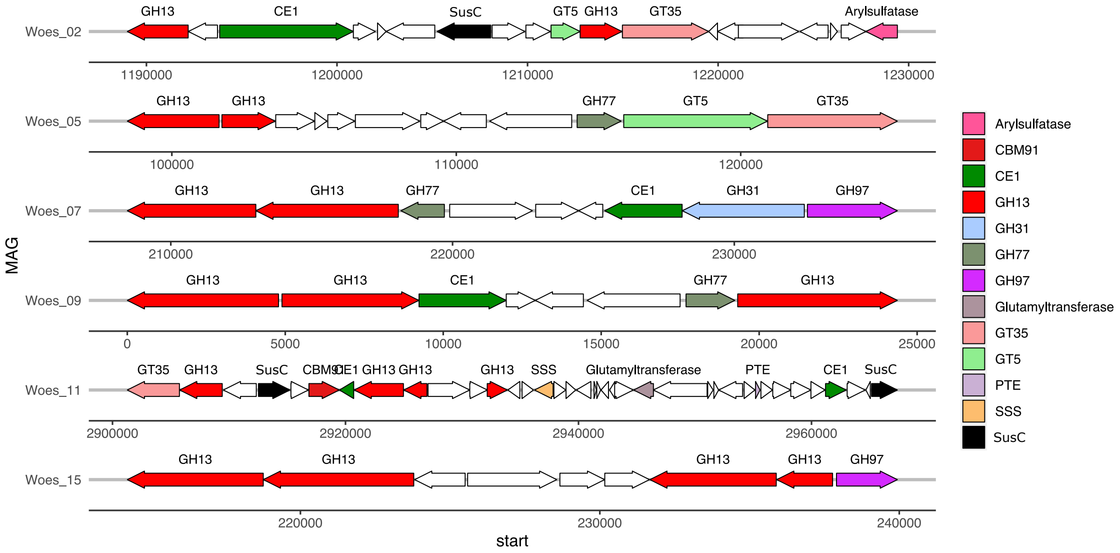

#### Figure S15: Genetic organization of the alginate PULs encoded by *Woeseiaceae* MAGs.

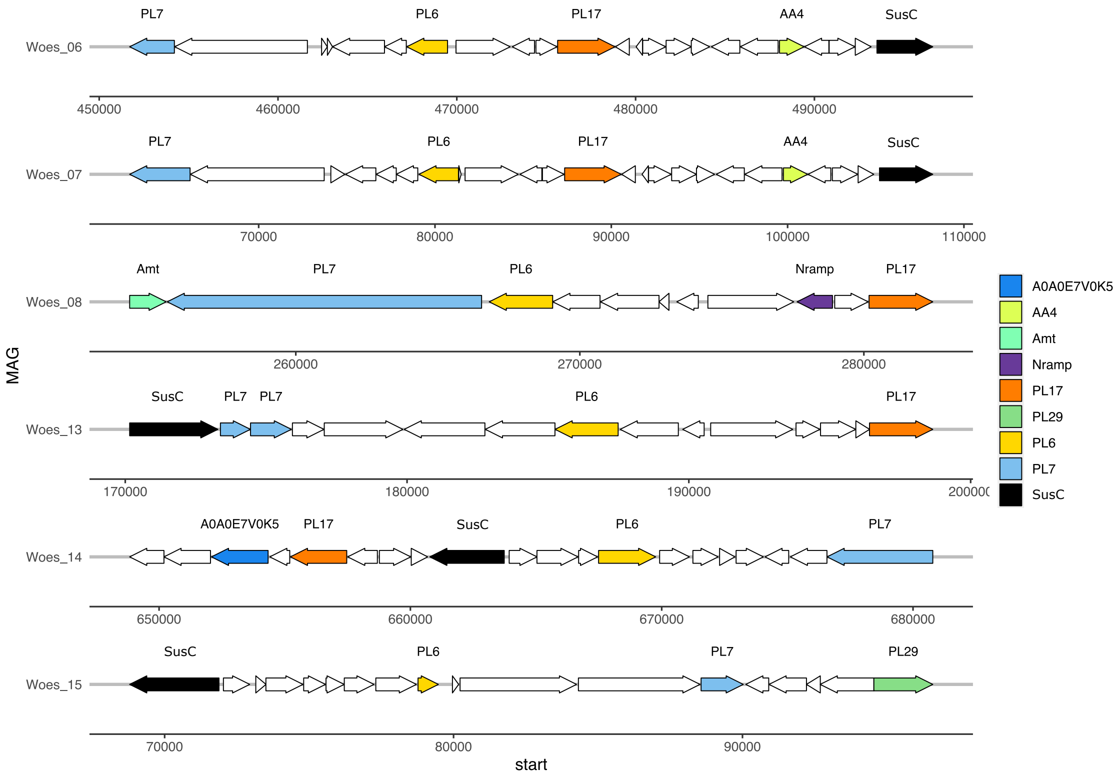

**SUPPLEMENTARY REFENCES**

[1] Miksch, S. *et al*. Bacterial communities in temperate and polar coastal sands are seasonally stable. *ISME Comm*. **1**, 29 (2021).
